# ATG5- and ATG7-mediated autophagy regulates pollen tube guidance and male fertility in *Arabidopsis thaliana*

**DOI:** 10.1101/2022.08.07.503073

**Authors:** He Yan, Xiaojuan Du, Mingkang Yang, Nianle Li, Xuequan Li, Zailue Ni, Wei Huang, Hong Wu, Lifeng Zhao, Hao Wang

**Affiliations:** Oollege of Life Sciences, South China Agricultural University, Guangzhou 510642, China; State Key Laboratory for Conservation and Utilization of Subtropical Agro-bioresources, South China Agricultural University, Guangzhou 510642, China

**Keywords:** ATG5, ATG7, autophagy, pollen germination, pollen tube, male fertility, sperm cell development, *Arabidopsis thaliana*

## Abstract

Autophagy functions as a crucial cellular scavenger by targeting cytoplasmic cargo to specific lysosome/vacuole for degradation. Autophagy-related (ATG) core proteins including ATG5 and ATG7 are evolutionarily conserved factors that are spatiotemporally orchestrated to regulate multiple processes of autophagy in yeast, mammalian and plant cells. However, autophagy is believed to be functionally dispensable in *Arabidopsis thaliana* since severe defects during plant growth, development and reproduction have not been observed in most of the *ATG* loss-of-function mutants, including *atg5* and *atg7,* under standard cultivation conditions. In this study, we report that autophagy does in fact play roles in regulating pollen tube growth guidance and male fertility in *Arabidopsis thaliana.* A detailed re-assessment of *atg5* and *atg7* mutants revealed greatly reduced autophagy activity in germinated pollens and the seed formation within siliques is partially abolished. Next, we demonstrated that both the pollen germination ratio and pollen tube length of the mutants decreased at the beginning of germination by time-lapse tracking of pollen germination *in vitro* and *in vivo.* Additionally, we observed occurrences of pollen tube twisting and stacking during their growth towards the ovules. Finally, we found abnormal pollen grains containing only a single sperm cell or an undivided generative nucleus. Collectively, these results indicate that ATG5- and ATG7-mediated autophagy is functionally involved in regulating pollen germination, tube growth guidance and sperm cell development in *Arabidopsis thaliana*.

## Introduction

Autophagy is an essential and conserved catabolic pathway in eukaryotic cells that functions as a cellular scavenger by targeting of cytosol and organelles to specific vacuoles or lysosomes for degradation (Liu and Bassham, 2012; Doherty and Baehrecke, 2018). Macroautophagy, the most prevalent type of autophagy and referred to as autophagy hereafter, serves as a housekeeping quality control mechanism that generally takes place at a basal and low level. When cells are challenged with biotic and/or abiotic stresses, the activity of autophagy will be significantly elevated to facilitate survival through the adverse conditions (Signorelli et al., 2019). In addition, autophagy plays versatile roles in regulating cell development, growth, reproduction and senescence (Ustun et al., 2017).

The key processes of autophagy consist of *de novo* formation and expansion of autophagosomal membranes, sequestration of unwanted cytoplasmic elements into the autophagosome and its subsequent fusion with a vacuole or lysosome. It is evident that ATG (autophagy-related) proteins play key roles during autophagy and have been identified in various species including yeasts, animals and plants (Klionsky et al., 2021). ATG proteins are spatiotemporally orchestrated to facilitate the biogenesis and functions of autophagosomes (Mizushima et al., 2011; Marshall and Vierstra, 2018). Moreover, approximately 18 *ATG* genes are found to be highly conserved across different species and therefore considered to be core *ATGs* for regulation of the autophagy machinery (Mizushima et al., 2011). In particular, ATG5 and ATG7 are considered to be particularly important because they are required for the autophagosome formation. ATG5 mediates the formation of the autophagosome precursor by conjugation with ATG12 to form a protein complex that catalyzes the chemical binding between cytosolic ATG8 and phosphatidylethanolamine on autophagosome membrane. ATG7 acts as an E1-like activating enzyme to facilitate the conjugation between ATG8-phosphatidylethanolamine and ATG12 during autophagosomal membrane expansion. ATG5 and ATG7 are often considered to be the two key factors for ATG8 lipidation which is a landmark event during autophagosome formation and autophagic flux (Liu and Bassham, 2012; Le Bars et al., 2014; Marshall and Vierstra, 2018; Yamaguchi et al., 2018; Klionsky et al., 2021). ATG5- and ATG7-knockout mice died within one day after birth due to the reduced concentration of cytosolic amino acids, a deleterious condition associated with autophagy deficiency (Akiko et al., 2004; Komatsu et al., 2005). Furthermore, ATG5- and ATG7-mediated autophagy has been found to regulate spermiogenesis and male fertility in mice (Wang et al., 2014; Huang et al., 2021). However, it is noteworthy that almost all of Arabidopsis *atg* mutants can survive well and accomplish their life cycle under ordinary cultivation conditions. Except for showing early senescence, Arabidopsis *atg5* and *atg7* mutant lines have no significant defects in growth and development and they successfully produce seeds. It is therefore believed that autophagy is not functionally required during Arabidopsis sexual reproduction (Marshall and Vierstra, 2018; Zhou et al., 2021).

Here, we argue that ATG5- and ATG7-mediated autophagy does indeed functionally participate in the process of sexual reproduction by regulating pollen tube growth and male fertility under nutrient-sufficient conditions in Arabidopsis. Despite previous reports that Arabidopsis *atg5* and *atg7* mutants can produce functional seeds and present few defects in pollen tube germination *in vitro,* we performed systematic and detailed investigations on the processes of pollen germination, pollen tube growth and male fertility of *atg5* and *atg7* null mutants. Our results suggest that autophagy is functionally involved in: i) maintenance of pollen tube growth and guidance to the ovules for fertilization; and ii) production of two functional sperm cells that are derived from the meiotic division of the generative nucleus during pollen development.

## Results

### ATG5 and ATG7 participate in autophagy in growing pollen tubes

To investigate whether autophagy plays a role in sexual reproduction in Arabidopsis, we employed *ATG5* and *ATG7* T-DNA insertion mutants to follow and compare flower development with that of wild type (WT). The schematic representation of the T-DNA insertion site within *atg5-1* and *atg7-2* is shown in Figure 1A. Quantitative RT-PCR (qRT-PCR) was performed to assess the gene expression levels of *ATG5* and *ATG7* in the *atg5-1* and *atg7-2* mutants respectively. *ATG5* and *ATG7* were barely detectable in *atg5-1* and *atg7-2* (Figure 1B). Moreover, we studied the gene expression profile of *ATG5* and *ATG7* in several different major cells and tissues of Arabidopsis by using the microarray data from Genevestigator (https://genevestigator.com/). Interestingly, we found that *ATG5* and *ATG7* are both enriched in pollens, sperm cells and stamens relative to other tissues in Arabidopsis (Supplemental Figure S1). Thereby, we performed western blotting to probe the level of autophagy activity in germinated pollens from both WT and the mutant Arabidopsis using an ATG8 antibody. ATG8 conjugation with phosphatidylethanolamine (ATG8-PE) is a reporter for the recruitment of cytosolic ATG8 onto the autophagosomal membrane for the subsequent biogenesis and functions of autophagosomes (Mizushima et al., 2011; Marshall and Vierstra, 2018). As shown in Figure 1C, ATG8-PE was barely detectable in *atg5-1* and *atg7-2* pollens when compared with the WT. This result indicates that autophagy is significantly impaired in *atg5-1* and *atg7-2* mutant pollens.

**Figure 1.**
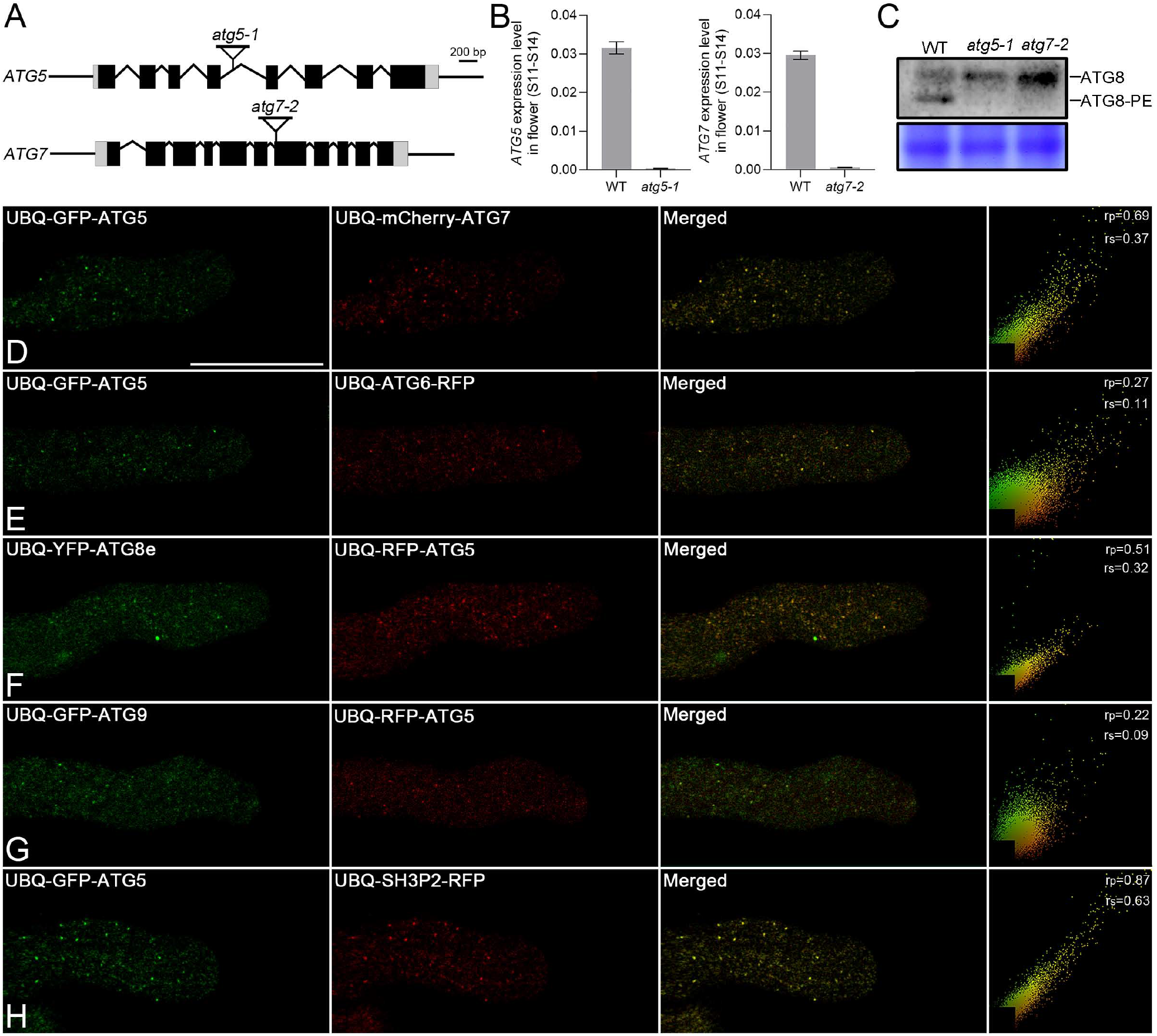
ATG5- and ATG7-mediated autophagy is involved in pollen tube growth in *Arabidopsis thaliana*. (**A**) Schematic illustration of the T-DNA insertion sites of *atg5-1* and *atg7-2* of *Arabidopsis thaliana*. (**B**) qRT-PCR analysis of expression of *ATG5* and *ATG7* in flowers (stage 11-14) of WT, *atg5-1* and *atg7-2. Actin2* was used as an internal control. Data are obtained from 3 independent experimental replicates (error bars ± SD) 0.01<*p< 0.05, **p < 0.01, (Student’s *t*-test). (**C**) Western blot detection of ATG8 and ATG8-PE in germinated pollen tubes of WT, *atg5-1* and *atg7-2* using ATG8 antibody. (**D-H**) Coexpression of GFP-ATG5 with (**D**) mCherry-ATG7 in growing tobacco pollen tubes. The punctate dots of ATG5 and ATG7 colocalize with one another and they mainly localize in the pollen tube shank. Furthermore, ATG5 were coexpressed with various autophagic markers including: (**E**) ATG6-RFP, (**F**) YFP-ATG8e, (**G**) GFP-ATG9 and (**H**) SH3P2-RFP in growing tobacco pollen tubes. Results of colocalization ratios are calculated by either Pearson correlation coefficients or as Spearman’s rank correlation coefficients with image J. The generated r values in the range −1 to 1, where 0 indicates no discernable correlation and +1 or −1 indicate strong positive or negative correlations, respectively. Scale bar in (**D**-**I**) = 25 μm.

To better understand the subcellular localization and spatial distribution of ATG5 and ATG7 in pollen tubes, we generated chimeric fluorescent fusion proteins of ATG5 and ATG7 and coexpressed them with several core autophagic proteins, including ATG6, ATG8e, ATG9 and SH3P2, in growing tobacco pollen tubes. ATG5 and ATG7 mainly colocalized with one another as shown in Figure 1D and Supplemental Movie S1. Additionally, ATG5 colocalized with ATG6, ATG8e, ATG9 and SH3P2 respectively (Figure 1E-H and Supplemental Movie S2-S5). These results suggest that ATG5 and ATG7 are involved in autophagy processes within growing pollen tube.

### The seed setting ratio is reduced in atg5 and atg7 Arabidopsis mutants

To further explore the regulatory functions of autophagy during the processes of sexual reproduction in Arabidopsis, we began by revisiting the overall phenotypes of plant growth and development during the reproductive stage.

Consistent with previous findings, no significant differences were observed for *atg5-1, atg7-2* and WT that had been cultivated for 35 and 45 d (Supplemental Figure S2). Furthermore, we compared the flower structures produced by *atg5-1*, *atg7-2* and WT. The top views of the flowers (stage 14) are shown in the first column of Supplemental Figure S3A-C. Additionally, the side view of stamen and pistil from stage 11 and 14 are shown and compared (Supplemental Figure S3A-C). Similarly to the overall growth and development phenotypes, no obvious developmental defects can be observed except that the length of pistils of both *atg5-1* and *atg7-2* are longer than that of the WT (Supplemental Figure S3A-C). Measurement and statistical analysis of *atg5-1*, *atg7-2* and WT pistil lengths are shown in Supplemental Figure S3D and E. This observation is consistent with and supported by the longer length of siliques produced by *atg5-1* and *atg7-2* compared that of the WT (Figure 2A-C). To test these results, we introduced *atg7-3,* which is another loss-of-function mutant generated by T-DNA insertion into *ATG7* as previously identified (Thompson et al., 2005). We again observed increased length of siliques (Figure 2D). Statistical analysis of the length of siliques from *atg5-1, atg7-2, atg7-3* and WT is shown in Figure 2E. Notably, a small portion of seed formation is abolished in the siliques of *atg5-1, atg7-2 and atg7-3* as indicated by the arrows (Figure 2F-I). Further statistical analysis by counting seed numbers within the siliques (>50) revealed that the seed setting ratios were actually reduced by ~20-25% in *atg5-1, atg7-2, atg7-3* mutants (Figure 2J).

**Figure 2.**
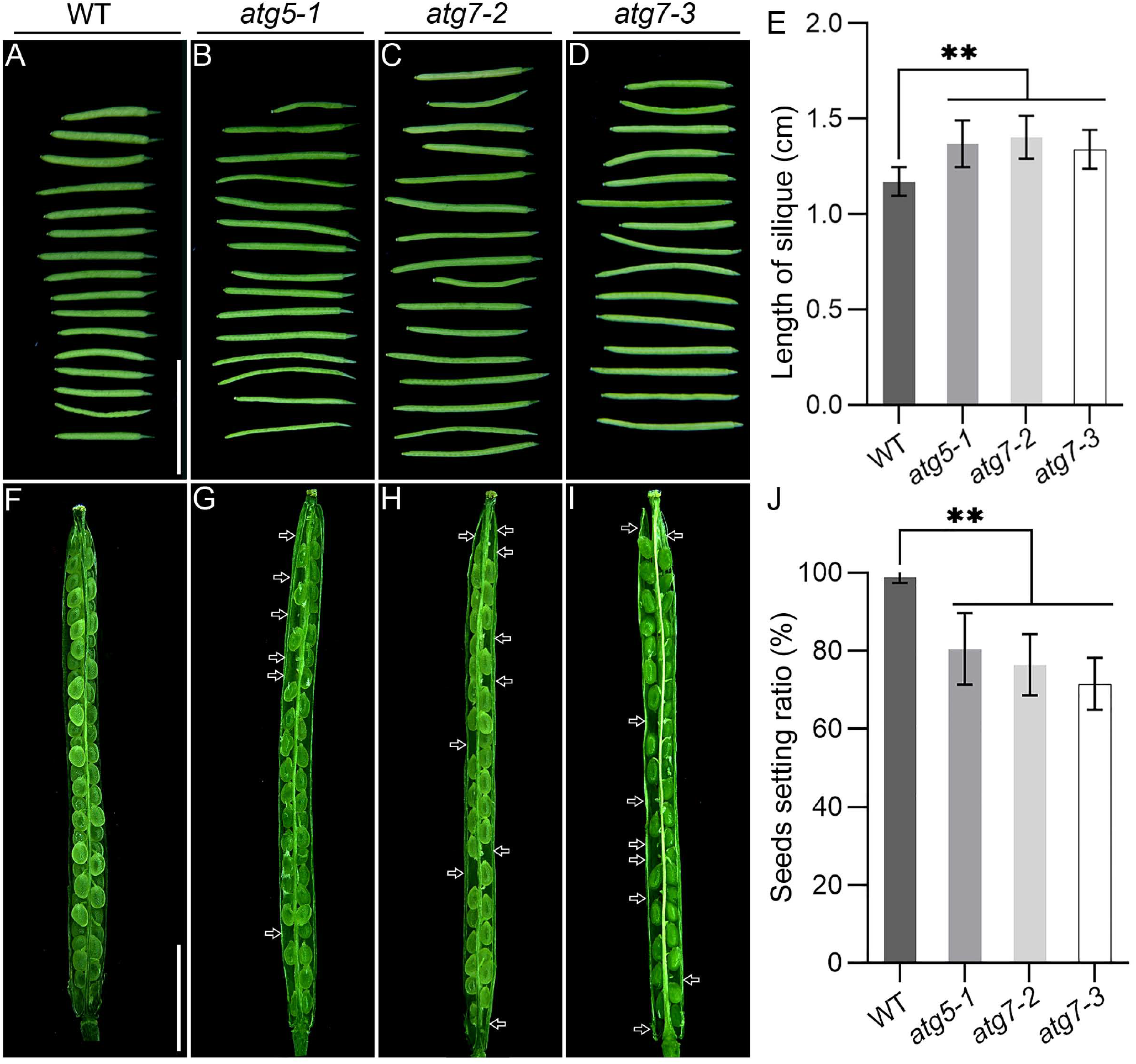
Seed formation is partially abolished in Arabidopsis *atg5* and *atg7* mutants. (**A-D**) Representative images of siliques harvested from one branch of inflorescence of Arabidopsis wild type (WT), *atg5-1, atg7-2* and *atg7-3.* **(E)** Statistical analysis of silique length of WT, *atg5-1, atg7-2* and *atg7-3* (error bars ± SD). 0.01<*p < 0.05, **p < 0.01 (Student’s *t*-test). More than 30 independent samples were calculated. (**F-I**) Representative images of seed setting in the siliques of WT, *atg5-1, atg7-2* and *atg7-3.* **(J)** Statistical calculation of seeds setting ratio in WT, *atg5-1, atg7-2* and *atg7-3* (error bars ± SD). Over 50 independent replicates from each sample were calculated. 0.01< *p < 0.05, **p < 0.01, (Student’s *t*-test). Scale bars in (**A-D**) = 1 mm, in (**F-I**) = 2 mm.

### ATG5- and ATG7-mediated autophagy regulates pollen germination, tube growth and male fertility in Arabidopsis thaliana

Previous independent studies found that both the pollen tube germination ratio and the pollen tube length of *atg5-1 and atg7-2* mutants were not distinct compared with WT after 10 h germination (Zhao et al., 2020). We revisited this analysis with a detailed time-lapse microscopy of *in vitro* pollen germination for 2, 4 and 6 h. Our results demonstrated that the pollen germination and pollen tube length of *atg5-1 and atg7-2* were significantly reduced when compared with WT at 2- and 4-h germinating timepoints (Figure 3A-C). In fact,-the pollen germination ratio of both *atg5-1 and atg7-2* reached only ~55% and ~80% in 2 and 4 h respectively, compared to nearly 80% and 90% for WT pollen at the relative timepoints (Figure 3D). Meanwhile, the length of pollen tubes of *atg5-1 and atg7-2* at 2-, 4- and 6-h germinated were obviously shorter compared with WT (Figure 3E).

**Figure 3.**
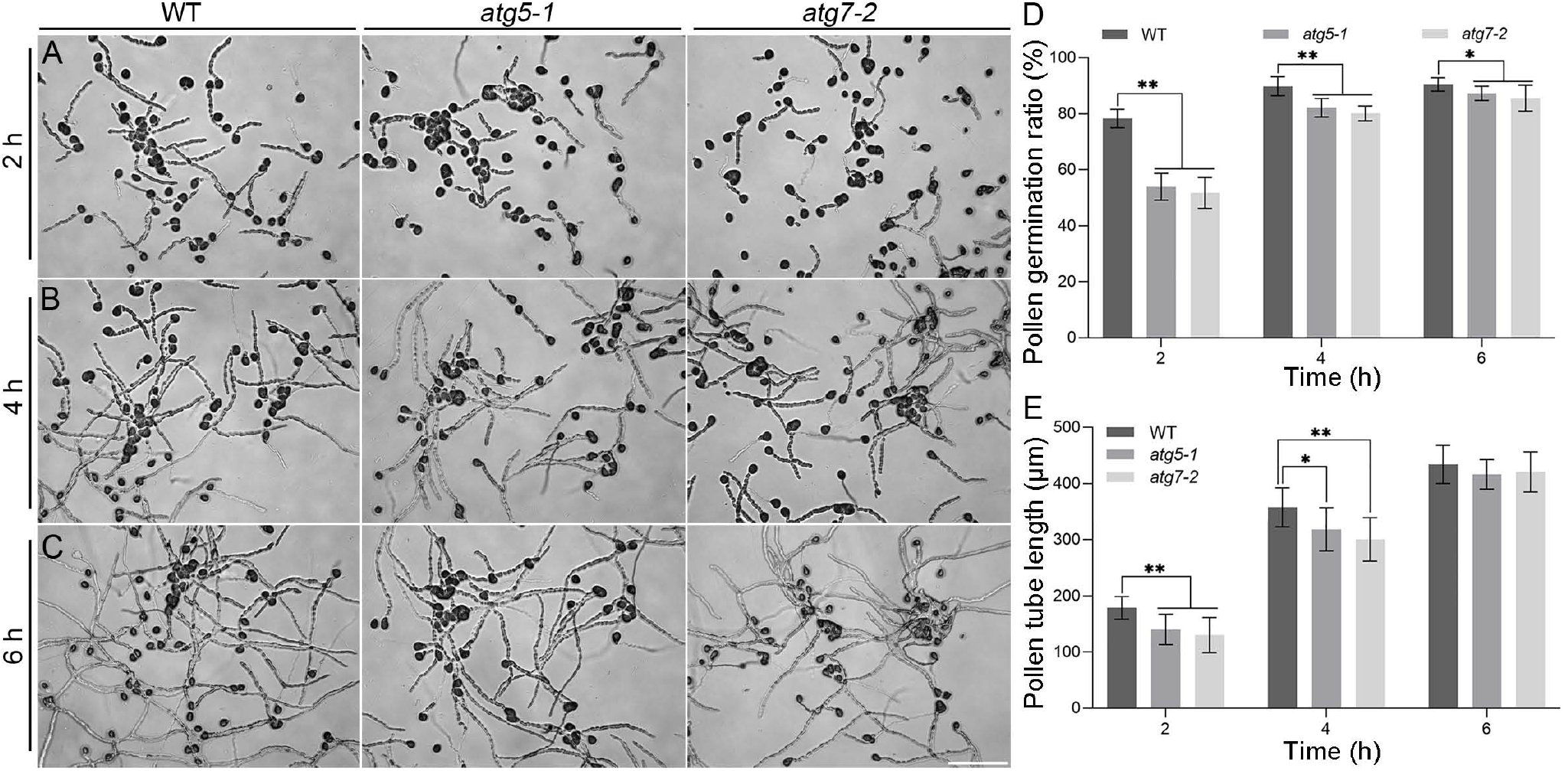
*In vitro* pollen tube germination and growth are reduced in Arabidopsis *atg5* and *atg7* mutants. (**A-C**) Representative images of *in vitro* Arabidopsis pollen tube germination and growth of WT, *atg5-1* and *atg7-2* for 2, 4 and 6 h respectively. Scale bar = 200 μm. (**D and E**) Statistical analysis of the *in vitro* pollen germination ratio and tube length of WT, *atg5-1* and *atg7-2* respectively. Over 500 pollens from each sample were calculated and measured (error bars ± SD). 0.01<*p < 0.05, **p < 0.01 (Student’s *t*-test).

We also conducted time-lapse microscopy to track pollen tube germination *in vivo* in WT and mutants of *atg5-1 and atg7-2* after 2-, 4-, 6- and 10-h pollination. The positions of maximal pollen tube growth within the styles are highlighted by the dashed lines (Figure 4A-D). The pollen tubes of *atg5-1 and atg7-2* were shorter than that of WT after 2-h pollination (Figure 4A). However, no significant differences of pollen tube growth can be observed after 4-, 6- and 10-h pollination (Figure 4B-D). Statistical analysis of the maximum lengths of pollen tubes within the styles of the WT and the mutants at different time points after pollination is shown in Figure 4E. In summary, pollen germination as assessed *in vitro* and *in vivo,* the pollen germination ratio and tube growth are all substantially reduced at the beginning of germination in *atg5-1 and atg7-2*.

**Figure 4.**
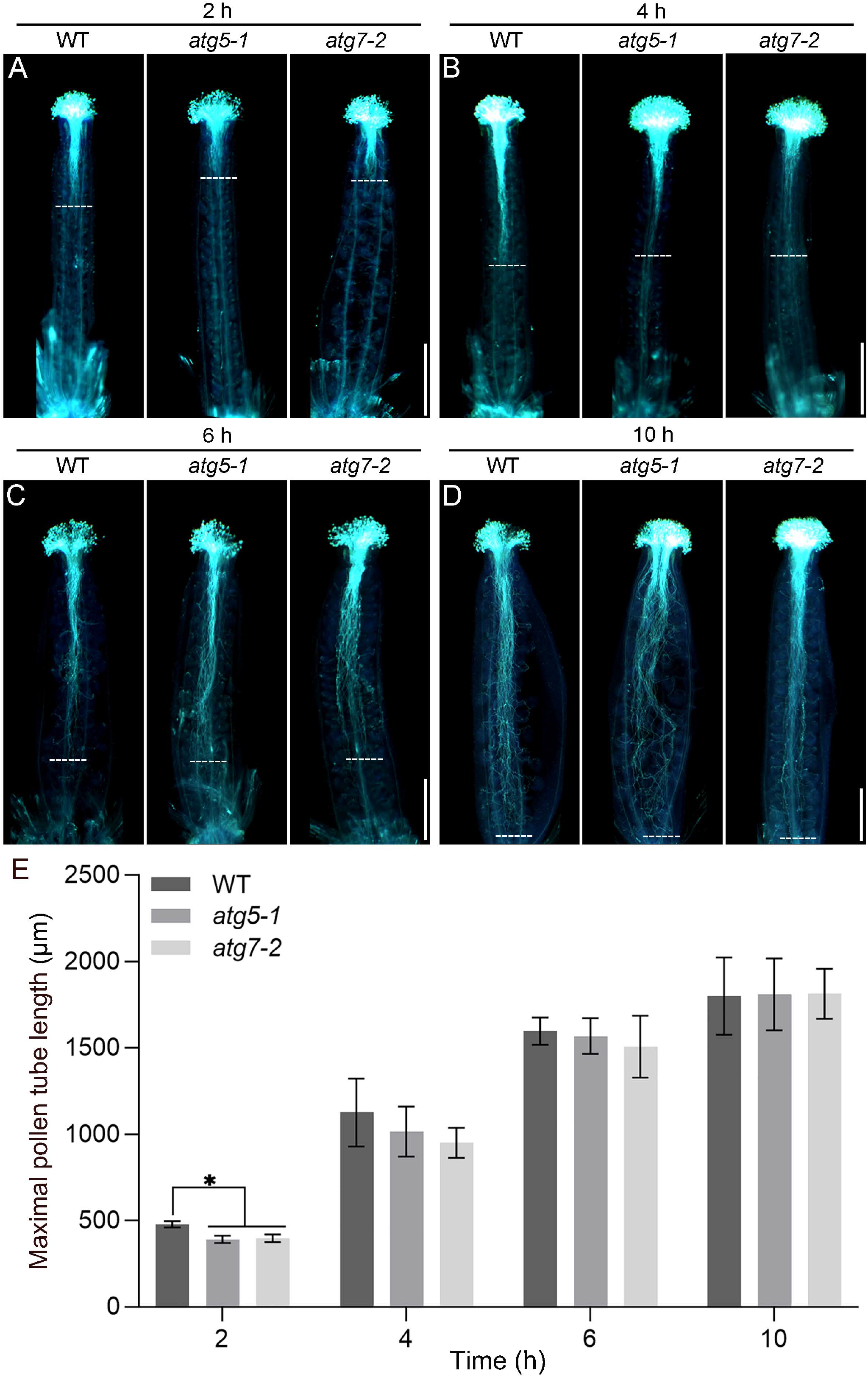
*In vivo* pollen tube germination and growth are reduced in Arabidopsis *atg5* and *atg7* mutants. (**A-D**) Representative images of *in vivo* pollen tube germination and growth of WT, *atg5-1* and *atg7-2* after pollination for 2, 4, 6 and 10 h respectively. Over 30 independent replicates of the individual samples were observed. Dashed lines indicate the farthest position that pollen tubes could reach into the styles. Scale bars = 400 μm. (**E**) Statistical calculation of the maximum pollen tube length in the styles of WT, *atg5-1* and *atg7-2* after 2-, 4-, 6- and10-h pollination. Over 50 replicates from each sample were measured (error bars ± SD). 0.01<*p < 0.05, **p < 0.01 (Student’s *t*-test).

To explore the underlying reason(s) accounting for the reduction of seed setting ratio for the mutants of *atg5-1 and atg7-2,* we investigated pollen tube guidance and targeting to the ovules. Pollen tubes in WT plants can grow generally straight and direct to the ovules for fertilization (Figure 5A and B). However, portions *atg5-1 and atg7-2* mutant pollen tubes grew twisted and accumulated outside the ovules as indicated by arrows in Figure 5D and F. In addition, we examined sperm cell development in mature pollens from WT, *atg5-1 and atg7-2* plants. After 4’,6-diamidino-2-phenylindole (DAPI) staining for 30 min, two well developed sperm cells can be clearly observed in WT pollen grains by confocal 3-D imaging (Figure 5G). Less than ~3% of WT pollens showed defects in sperm cell development (Figure 5J). In contrast, ~15% of mature pollens from *atg5-1 and atg7-2* strains contained only one sperm cell/undivided generative nucleus as indicated by arrows in Figure 5H and I. Statistical calculations are shown in Figure 5J.

**Figure 5.**
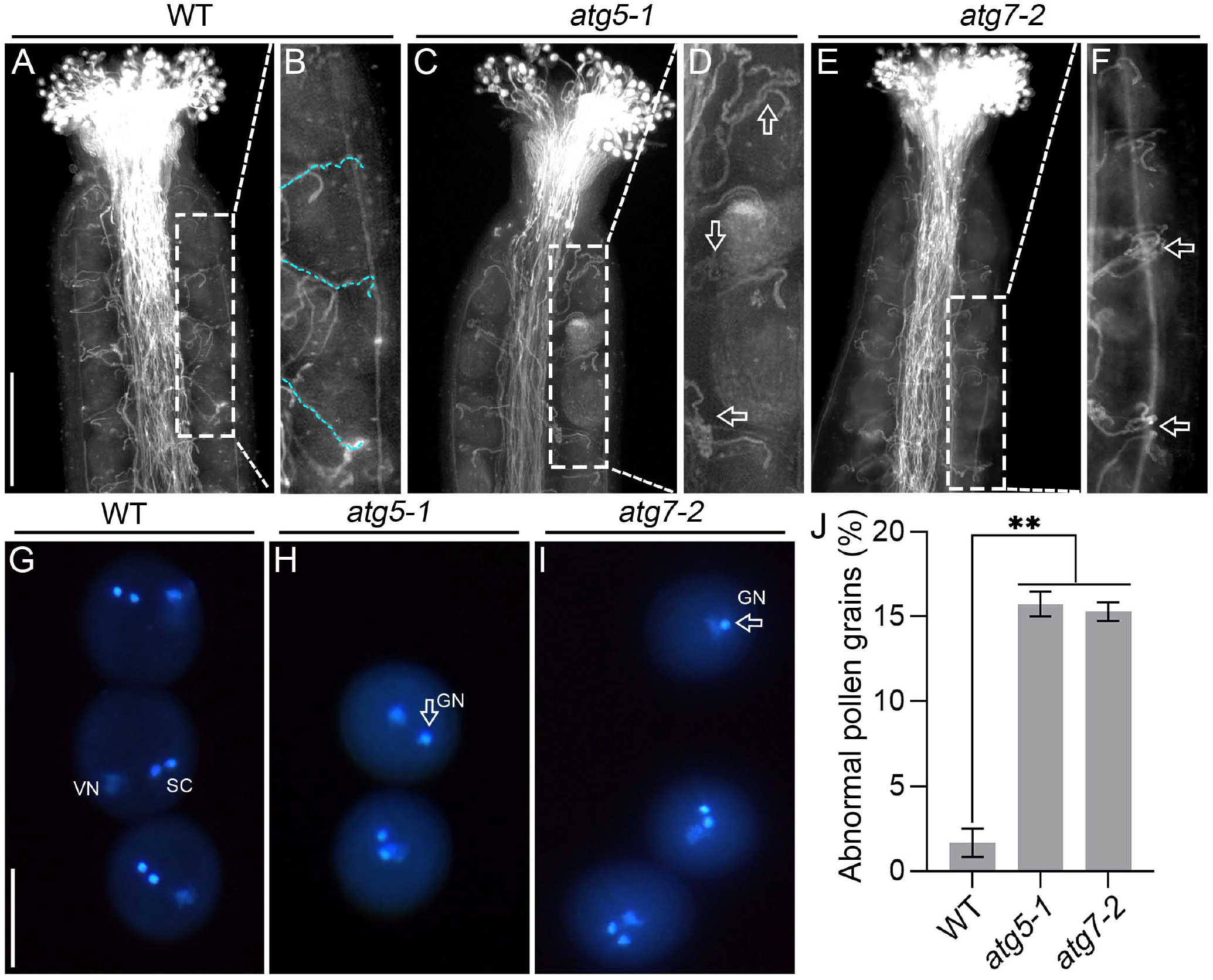
Defects in pollen tube growth guidance and sperm cell formation in *atg5-1* and *atg7-2* of *Arabidopsis thaliana*. (**A**) Pollen tube growth guidance to the ovules in WT Arabidopsis. (**B**) Enlarged image derived from the selected area in (**A**) shows the detail of the pollen tubes growth guidance and targeting to the ovules. (**C**) Pollen tube growth guidance to the ovules in *atg5-1.* (**D**) Enlarged image derived from the selected area in (**C**) demonstrates the occurrence of twisted pollen tubes that accumulated outside the ovules as indicated by the arrow. (**E**) Pollen tube growth guidance to the ovules in *atg7-2.* (**F**) Enlarged image derived from the selected area in (**E**) shows the occurrence of twisted pollen tubes that accumulated outside the ovules as indicated by the arrow. Scale bar in (**A**), (**C**) and (**E**) = 100 μm. (**G-I**) DAPI staining of nucleus in mature pollen grains of WT, *atg5-1* and *atg7-2* respectively. (**G**) A representative image shows that WT individual pollens containing two sperm cells and a vegetative nucleus. (**H-I**) Representative images of mature pollens from *atg5-1* and *atg7-2.* Some of the mutant pollen grains only contain a single sperm cell/undivided generative nucleus as indicated by the arrow in (**H**) and (**I**) respectively. Scale bar in (**G-I**) = 10 μm. (**J**) Statistical analysis of abnormal pollen ratio in the mature pollens of WT, *atg5-1* and *atg7-2.* Over 200 mature pollens from each sample were calculated (error bars ± SD). 0.01<*p < 0.05, **p < 0.01 (Student’s *t*-test).

## Discussion

Recent evidence has shown that autophagy is involved in multiple steps of sexual reproduction, depending on plant species, including male gametophyte development and maturation, self-incompatible pollen rejection and pollen germination (Harrison-Lowe and Olsen, 2008; Kurusu et al., 2014; Li et al., 2020; Norizuki et al., 2020; Zhao et al., 2020; Zhou et al., 2021; Macgregor et al., 2022). Nevertheless, the role(s) of canonical autophagy during sexual reproduction in *Arabidopsis thaliana* is still an enigma since almost all of the autophagy-defective Arabidopsis mutants such as *atg2*, *atg5* and *atg7* exhibit normal life cycles without any deleterious phenotypes during fertilization, embryogenesis and seed formation. Plants with these mutations are comparable to WT Arabidopsis with regards to progression and completion of their life cycle (Marshall and Vierstra, 2018; Zhou et al., 2021). On the other hand, Arabidopsis plants with the *ATG6* loss-of-function mutation show substantial defects in pollen germination (Qin et al., 2007; Harrison-Lowe and Olsen, 2008). However, ATG6 is a key component of the phosphoinositide 3-kinase (PtdIns3K) complex and therefore involved in many intracellular processes including endocytosis and endomembrane trafficking (Liu and Bassham, 2012; Marshall and Vierstra, 2018). It is currently not possible to rule out these various activities as they affect pollen germination. The exact roles of ATG6 in regulating pollen germination *via* autophagy remain to be further studied and characterized. Due to the lack of substantial, definable phenotypes of these ATG mutants, autophagy is widely considered to be functionally irrelevant during reproduction in *Arabidopsis thaliana.* Despite these observations, it is curious and noteworthy that several core *ATG*s, including *ATG5, ATG7 and ATG8h,* are highly enriched in anthers or pollens compared with other tissues based on expression profile analyses (Supplementary Figure S1) (Zhou et al., 2021). Additionally, western blot analysis of ATG8-PE in germinated Arabidopsis pollens showed substantially reduced autophagic activity in *atg5-1* and *atg7-2* mutants compared with WT (Figure 1). These results suggest a role for autophagy in regulating functions of male gametophytes in Arabidopsis.

In this study, we argue that ATG5- and ATG7-mediated autophagy indeed participates in regulating the reproduction of *Arabidopsis thaliana* by controlling pollen tube growth guidance and male fertility. This conclusion is supported by several lines of evidence: i) detailed investigation on the seed formation in siliques of *atg5-1, atg7-2* and *atg7-3* revealed that ~15-20% reduction of seed setting compared with WT (Figure 2). This result could be overlooked since the seed formation defect in *atg5-1, atg7-2* and *atg7-3* is not severe; ii) the pollen germination ratio and tube growth of *atg5-1* and *atg7-2* were reduced, especially at the beginning of germination, as demonstrated by time-lapse tracking of pollen tube germination (Figure 3 and 4). Previous studies allowed the pollen from *atg5-1* and *atg7-2* strains to germinate for 10 h or more, which could lead to the overgrowth of pollen tubes and, therefore, obscure the phenotype (Liu and Bassham, 2012; Zhou et al., 2021); iii) more detailed studies of pollen tube guidance and targeting to ovules showed that a portion of *atg5-1* and *atg7-2* pollen tubes were deformed. They became twisted and-accumulated outside of the ovules rather than growing straight to the ovules for fertilization as observed for WT pollen tubes (Figure 5). Although it remains to be determined whether these abnormal pollen tubes could eventually achieve fertilization, it is evident that autophagy is functionally involved in pollen tube guidance and targeting during fertilization in *Arabidopsis thaliana*; iv) ~15% of the mature pollens of *atg5-1* and *atg7-2* contain only a single sperm cell or undivided generative nucleus. This result likely accounts for the aberrant seed formation within the siliques of the mutants (Figure 2 and 5).

Emerging studies show that ATG5- and ATG7-mediated autophagy participates in the clearance of cytoplasm in the process of sperm cell differentiation in lower plants that generate motile sperm cells for fertilization (Sanchez-Vera et al., 2017). In addition, the loss-of-function mutants of OsATG7 and OsATG9 cause male sterility in rice (Kurusu et al., 2014). Far fewer autophagosome-like structures were observed in the tapetum of *Osatg7* rice (Kurusu et al., 2014). Even though the underlying cellular mechanisms in the tapetum require further investigation, this observation indicates that autophagy is required for the development of male gametes in rice. Consistently, our results demonstrate that ATG5- and ATG7-mediated autophagy functionally participates in regulating pollen tube growth guidance and male fertility under standard cultivation conditions in *Arabidopsis thaliana*. In fact, it is shown that canonical ATG5- and ATG7-mediated autophagy is required for normal sperm formation and male fertility in animals. For example, ATG5 and ATG7 were shown to induce autophagy by integrating multiple signals to maintain normal developmental processes of male fertility in mice (Wang et al., 2014; Liu et al., 2016; Huang et al., 2021). ATG5-mediated autophagy controls various aspects of spermiogenesis including spermatid development, sperm individualization and sperm fertility (Huang et al., 2021). Furthermore, the depletion of ATG7 disrupted autophagic flux in germ cells and resulted in irregular or nearly round-headed spermatozoa (Wang et al., 2014).

The state of research focused on elucidating the roles of autophagy and their mechanistic details in plant reproduction is still in its infancy. Characterization of these processes will be of great interest and importance towards understanding the evolutionary conservation and diversification of autophagy in regulating male fertility in animals compared with that of plants. One open and interesting question to address is how ATG5- and ATG7-mediated autophagy regulates the formation of two sperm cells that are derived from meiosis of the generative nucleus in *Arabidopsis thaliana*.

## Materials and Methods

### Plant materials and growth conditions

*Arabidopsis thaliana* wild type (Col-0) and *atg5-1, atg7-2 and atg7-3* T-DNA insertional mutants in the ecotype Col background were used in this study. Arabidopsis seeds were surface sterilized and germinated on Murashige and Skoog (MS) medium containing 0.8% (w/v) agar and 3% (w/v) sucrose, pH 5.7. Seedlings were cultured in a plant growth chamber at 22°C under a 16-light and 8-h dark cycle prior to being transferred into soil in a plant growth room at 22°C under a 16-h light and 8-h dark cycle. The light intensity was ~150 μmol/m^2^/s.

### Transient expression of chimeric fluorescent fusion proteins in tobacco pollen tubes by particle bombardment

In brief, anthers were harvested freshly from ~20 tobacco flowers and transferred to 20 ml tobacco pollen germination medium. Pollens were released into the medium from anthers by vigorous agitation with a vortex for 2 min. Subsequent preparation of pollen grains and transient expression of proteins in growing pollen tubes *via* biolistic bombardment were performed as previously described (Wang and Jiang, 2011; Zhong et al., 2017).

### Confocal laser scanning microscopy imaging and colocalization calculations

In general, confocal fluorescent images were collected by using a Leica TCS SP8 system with the following settings: 63x water immersion objective, 2x zoom, 750 gain, 0 background, 0.168 μm pixel size and photomultiplier tubes (PMTs) detector. A confocal dish (NEST, cat no. 801002) with a cover slide at the bottom was used for live-cell imaging of pollen tubes to minimize potential mechanical damages caused by the conventional sandwich method of a sample between glass and cover slides. Colocalization analysis between two fluorescent proteins was calculated using ImageJ software (https://imagej.nih.gov/ij/) with the Pearson-Spearman correlation (PSC) colocalization plug-in (French et al., 2008). Results were presented either as Pearson correlation coefficients or as Spearman’s rank correlation coefficients, both of which produce r values in the range (−1 to 1), where 0 indicates no discernable correlation and +1 or −1 indicate strong positive or negative correlations, respectively.

### Total protein extraction and western blotting

Total proteins were extracted from freshly opened Arabidopsis flowers. The protocol of protein extraction followed a previously optimized method for better detection of ATG8-lipid adduct (Chung et al., 2010). Equal amounts of proteins were loaded onto a 15% SDS-PAGE gel with 6 M urea. Thereafter, proteins were transferred onto a polyvinylidene fluoride (PVDF) membrane. ATG8-PE and free ATG8 were immunodetected by anti-ATG8 antibody (Abcam, cat no. ab77003).

### qRT-PCR

Total RNA was isolated from freshly opened Arabidopsis flowers (S14) (Vazyme, cat no. RC401-01). The first strand for each cDNA was synthesized by HiScript II reverse transcriptase (Vazyme, cat no. R312-01). Real-time fluorescence qRT-PCR was performed on Bio-Rad CFX Connect Real-Time PCR Detection System. At least three independent replicates were performed. *Actin2* was used as an internal control.

### In vitro and in vivo Arabidopsis pollen germination

For *in vitro* pollen germination: mature pollens at the specific developmental stage were collected as previously described (Mori et al., 2006; Zhong et al., 2017). Mature Arabidopsis pollen grains were transferred from anthers by spreading them on Arabidopsis solid pollen germination medium containing 0.01% boric acid, 1 mM Ca(NO_3_)2, 1 mM MgSO_4_, 5 mM CaCl_2_, 18% (w/v) sucrose and 0.9% agarose at pH 7.0 on an standard glass microscopy slide. The slide was then incubated in a wet chamber box at 28°C and germinated for 2, 4 and 6 h prior to imaging.

For *in vivo* pollen germination: Arabidopsis flowers (stage 12) were castrated and used as female recipients. Pollen grains were selected from stage 15 flowers and gently spread onto the surface of stigma of the female recipients for *in vivo* pollen tube germination. After 2-, 4-, 6-, 10- and 16-h pollination, the female tissues were fixed with acetic acid/ethanol 1:3 (v/v) solution for 2 h at RT. Then, the samples were rehydrated by a series of incubations in 70%, 50% and 30% ethanol for 10 min each, and rinsed with ddH_2_O for 10 min at RT. They were subsequently immersed in 8 M NaOH overnight at RT and then washed for 3 times with ddH_2_O. Finally, the samples were stained by aniline blue solution containing 0.2% (w/v) aniline blue and 0.1 M K_3_PO_4_ for 2-3 h at RT. Confocal laser scanning microscopy imaging was then immediately performed using a TCS SP8 confocal microscope (Leica).

### DAPI staining

Mature pollen grains were released from freshly opened flowers by vortexing them in Arabidopsis pollen germination medium containing 0.01% boric acid, 1 mM Ca(NO_3_)_2_, 1 mM MgSO_4_, 5 mM CaCl_2_ and 18% (w/v) sucrose at pH 7.0. The flower petals and anthers were removed with forceps. Arabidopsis pollens were concentrated by centrifugation at 800 g for 1 min at RT. After discarding the supernatant, the pellet of pollens was gently and fully resuspended with 200 μl DAPI staining solution containing 0.25 M sodium phosphate (pH 7.0), 0.25 mM EDTA, 0.025% Triton X-100 and 0.1 μg/ml DAPI (Sangon Biotech, cat no. E607303) and incubated in the dark for 30-45 min. Confocal laser scanning microscopy imaging was then immediately performed using a TCS SP8 confocal microscope (Leica).

### Accession numbers

The Arabidopsis Genome Initiative locus identifiers for the genes mentioned in this article are *ATG5* (AT5G17290), *ATG6* (AT3G61710), *ATG7* (AT5G45900), *ATG8e* (AT2G45170), *ATG9* (AT2G31260), *SH3P2* (AT4G34660).

## Supplemental Data

**Supplemental Figure S1.** Gene expression profile analysis of *ATG5* and *ATG7* genes in several Arabidopsis tissues.

**Supplemental Figure S2.** The representative overview images of WT, *atg5-1* and *atg7-2* of Arabidopsis thaliana during the reproduction period.

**Supplemental Figure S3.** Phenotypes of WT, *atg5-1* and *atg7-2* flowers.

**Supplemental Movie S1.** Co-expression of GFP-ATG5 with mCherry-ATG7 in growing tobacco pollen tubes.

**Supplemental Movie S2.** Co-expression of GFP-ATG5 with ATG6-RFP in growing tobacco pollen tubes.

**Supplemental Movie S3.** Co-expression of RFP-ATG5 with YFP-ATG8e in growing tobacco pollen tubes.

**Supplemental Movie S4.** Co-expression of RFP-ATG5 with GFP-ATG9 in growing tobacco pollen tubes.

**Supplemental Movie S5.** Co-expression of GFP-ATG5 with SH3P2-RFP in growing tobacco pollen tubes.

## Acknowledgments

We apologize to those whose work could not be cited because of space restrictions. We would like to thank the members of Wang laboratory for stimulating discussions. We also thank Dr. Andrew Loria for help with editing and Dr. Faqiang Li (South China Agricultural University) for providing *atg5-1, atg7-2* and *atg7-3* mutant seeds (*A. thaliana* Col-0).

