## Supplemental Figures for "ATG5- and ATG7-mediated autophagy regulates pollen tube guidance and male fertility in *Arabidopsis thaliana*"

### Title Page

**Affinities:**

[1]College of Life Sciences, South China Agricultural University, Guangzhou 510642, China.

[2]State Key Laboratory for Conservation and Utilization of Subtropical Agro-bioresources, South China Agricultural University, Guangzhou 510642, China.

[3]These authors contributed equally to the article.

*Senior author.

***Corresponding author:**

Hao Wang, Ph.D., Professor

**Keywords:** ATG5, ATG7, autophagy, pollen germination, pollen tube, male fertility, sperm cell development, *Arabidopsis thaliana*

**Running title:** Autophagy regulates Arabidopsis male fertility

**Abstract**

Autophagy functions as a crucial cellular scavenger by targeting cytoplasmic cargo to specific lysosome/vacuole for degradation. Autophagy-related (ATG) core proteins including ATG5 and ATG7 are evolutionarily conserved factors that are spatiotemporally orchestrated to regulate multiple processes of autophagy in yeast, mammalian and plant cells. However, autophagy is believed to be functionally dispensable in *Arabidopsis thaliana* since severe defects during plant growth, development and reproduction have not been observed in most of the *ATG* loss-of-function mutants, including *atg5* and *atg7*, under standard cultivation conditions. In this study, we report that autophagy does in fact play roles in regulating pollen tube growth guidance and male fertility in *Arabidopsis thaliana*. A detailed re-assessment of *atg5* and *atg7* mutants revealed greatly reduced autophagy activity in germinated pollens and the seed formation within siliques is partially abolished. Next, we demonstrated that both the pollen germination ratio and pollen tube length of the mutants decreased at the beginning of germination by time-lapse tracking of pollen germination *in vitro* and *in vivo.* Additionally, we observed occurrences of pollen tube twisting and stacking during their growth towards the ovules. Finally, we found abnormal pollen grains containing only a single sperm cell or an undivided generative nucleus. Collectively, these results indicate that ATG5- and ATG7-mediated autophagy is functionally involved in regulating pollen germination, tube growth guidance and sperm cell development in *Arabidopsis thaliana*.

**Supplemental Figures**

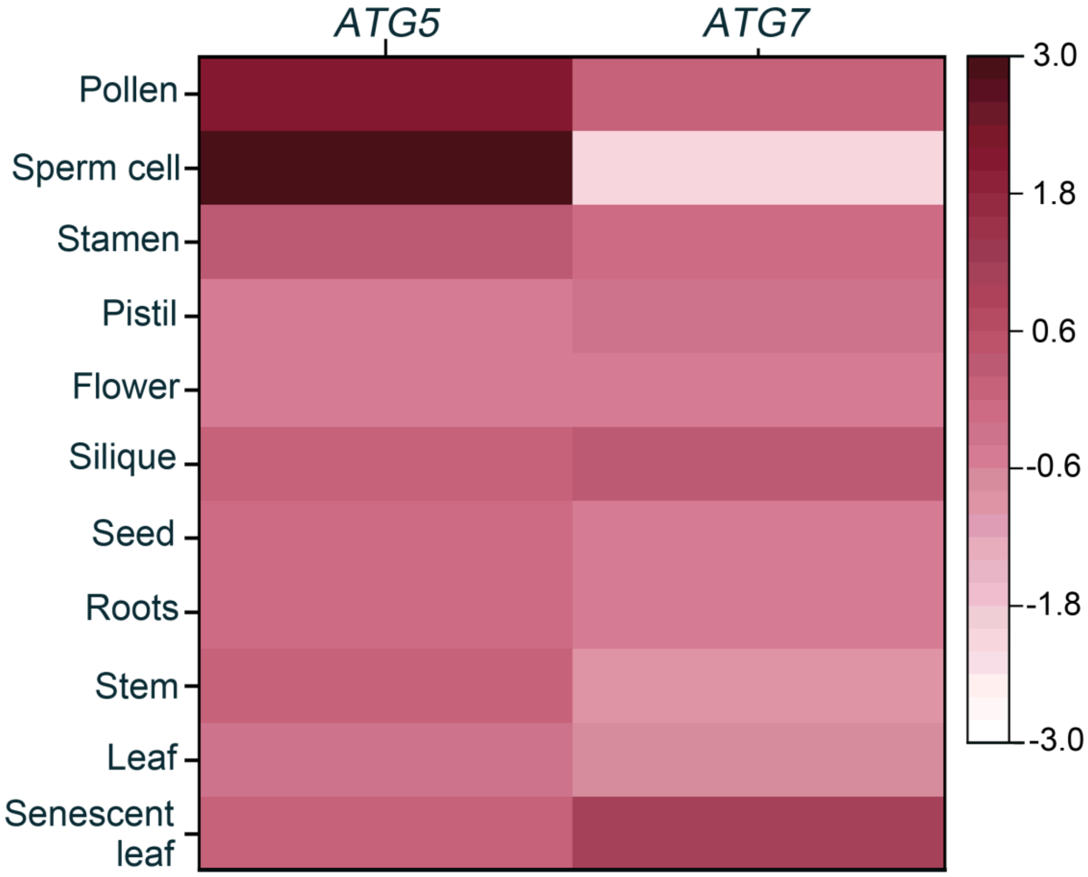


**Supplemental Figure S1.** Gene expression profile analysis of *ATG5* and *ATG7* genes in several Arabidopsis tissues.

The expression profile for *ATG5* and *ATG7* were obtained from Genevestigator (https://genevestigator.com/gv/). The heatmap image was created by HEATMAPPER as described previously (Babicki et al., 2016). The dark red box represents higher gene expression level, whereas the light red box represents lower gene expression level. The scale bar represents the raw z-score.

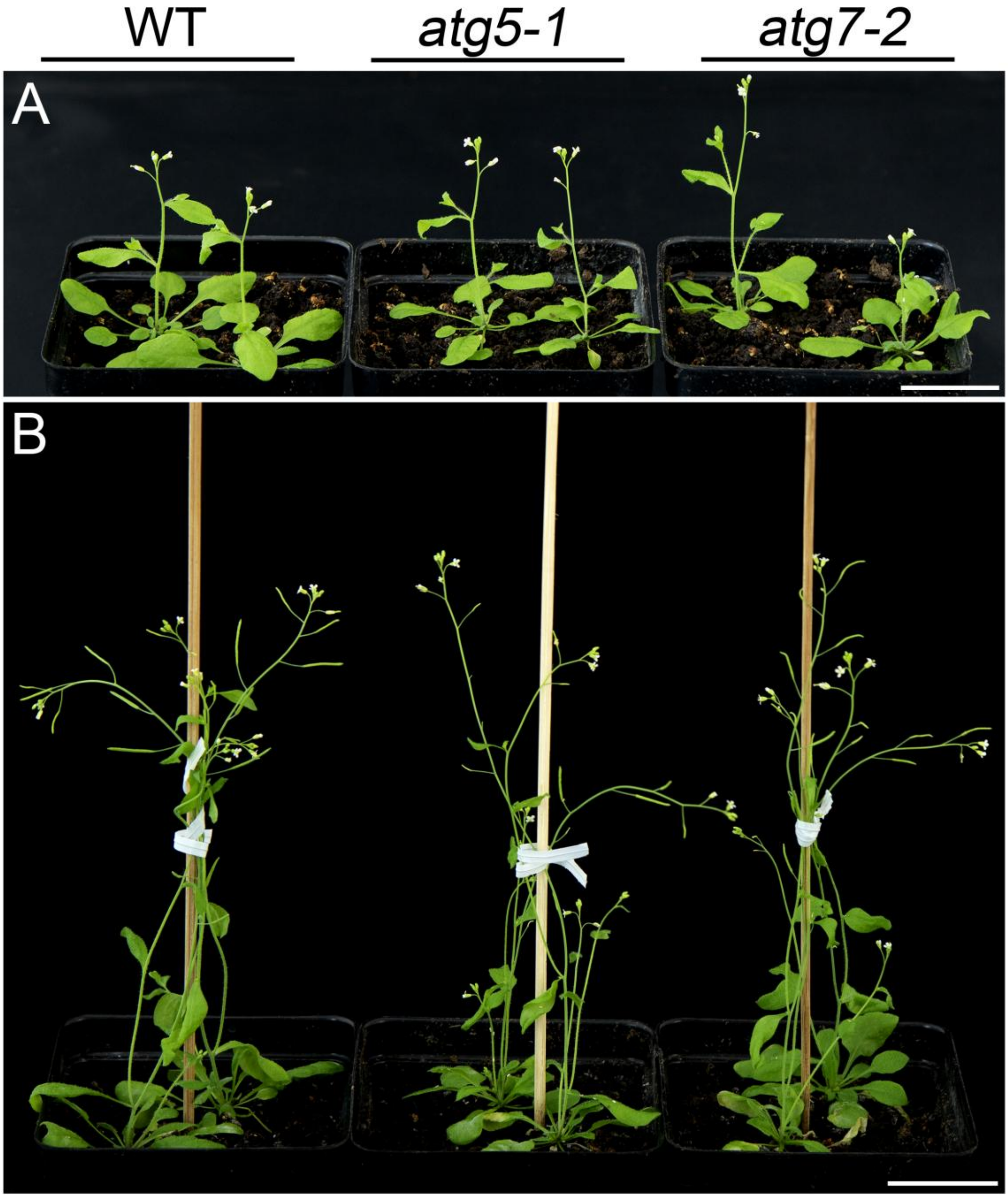


**Supplemental Figure S2.** The representative overview images of WT, *atg5-1* and atg*7-2* of *Arabidopsis thaliana* during the reproduction period.

The overview of WT, *atg5-1* and atg*7-2* *Arabidopsis thaliana* at the reproductive stage cultivated for 35 days (**A**) and 45 days (**B**). Scale bar = 3 cm.

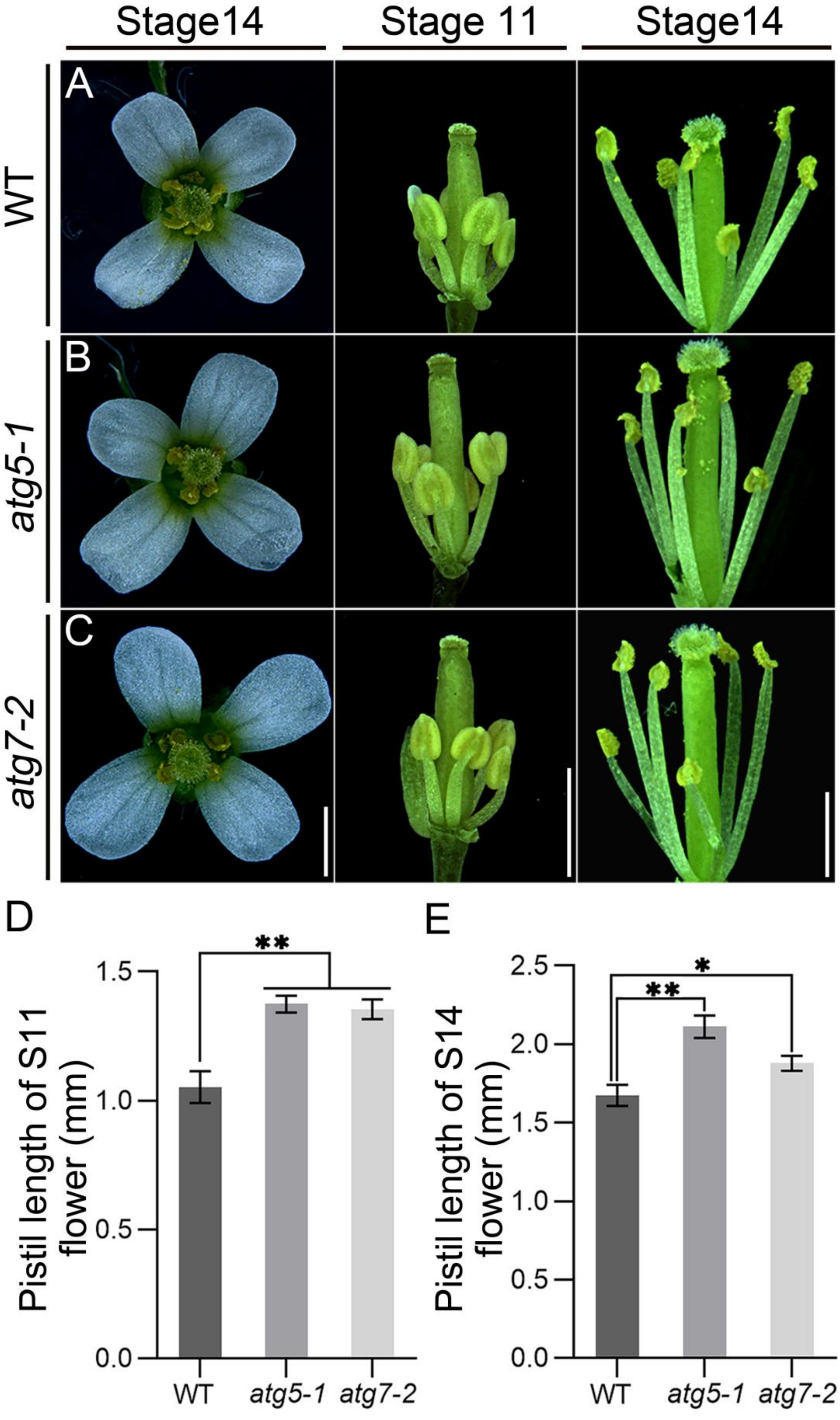
Stage14
Stage 11
Stage14
A
B
C
WT
atg5-1
atg7-2
D
E
**
Pistil length of S11 flower (mm)
1.5
1.0
0.5
0.0
WT
atg5-1
atg7-2
*
**
Pistil length of S14 flower (mm)
2.5
2.0
1.5
1.0
0.5
0.0
WT
atg5-1
atg7-2

**Supplemental Figure S3.** Phenotypes of WT, *atg5-1* and *atg7-2* flowers.

(**A-C**) Representative images of the flowers of WT, *atg5-1* and atg*7-2* *Arabidopsis thaliana*. The top overview the stage 14 flowers and the side view of stage 11 and 14 flowers whose sepals and petals were removed to reveal elongated gynoecium. No significant differences of androecium were observed between the WT and the *atg5-1* and *atg7-2* mutants. Scale bars =1 mm.

(**D** and **E**) Statistical analysis of the pistil length of WT, *atg5-1* and atg*7-2*. Over 50 independent replicates of each sample were measured (error bars ± SD). $0.01 < *p < 0.05$, $**p < 0.01$ (Student's *t*-test).
